# Rab27a modulation reverses hemorrhage-induced neutrophil dysfunction in response to secondary respiratory challenge

**DOI:** 10.64898/2026.11.10.765693

**Authors:** Anandita Basu, Anirudh Udutha, Sean Park, Lillian I. Levin, Melanie A. Martinsen, Ludimira S. Ribeiro, Craig T. Lefort

## Abstract

Patients recovering from hemorrhage are susceptible to secondary respiratory infections, owing to dysregulated innate immune responses. Using a two-hit mouse model of hemorrhage followed by a secondary respiratory challenge with *Staphylococcus aureus* or lipopolysaccharide (LPS), we investigated the role of Rab27a in the neutrophil response to pulmonary inflammation and host defense. Hemorrhage followed by *S. aureus* infection increased surface CD63 expression on pulmonary neutrophils, indicating azurophilic granule release, and was accompanied by reduced neutrophil entry into the airspaces of the lung. Using HoxB8-conditional neutrophil progenitor (NP) transplantation to neutrophil chimeric mice, we observe that Rab27a-deficient neutrophils have enhanced recruitment into the extravascular pulmonary compartment compared with wild-type (WT) neutrophils. Furthermore, pharmacological intervention with Nexinhib20, a small-molecule inhibitor of Rab27a-JFC1 interactions, attenuated systemic and pulmonary inflammatory responses in the two-hit model, including a reduction in circulating myeloperoxidase (MPO) levels. Nexinhib20 also enhanced neutrophil-mediated *S. aureus* killing that was accompanied by increased intracellular MPO levels in neutrophils. Collectively, these findings implicate Rab27a-associated functions in the regulation of neutrophil trafficking and inflammatory responses following hemorrhage and secondary respiratory challenge, highlighting the potential of pharmacological targeting of Rab27a to attenuate inflammation without compromising antimicrobial host defense.

## Introduction

Hemorrhage and its sequelae are responsible for more than a third of deaths due to injury in the United States^1^. Due to the excessive loss of blood, patients with hemorrhage have an increased susceptibility to infections owing to a cascade of events leading to immunosupression^2^. Such immunosuppression could finally lead to multi-organ failure of which acute respiratory distress syndrome (ARDS) is one of the most common occurrences^3^. Pneumonia remains the most frequent cause of ARDS^4^. Methicillin-resistant *S. aureus* is a detrimental pathogen contributing to pulmonary infections in the critically ill^5^ and is responsible for 20-40% of all nosocomial pneumonias^6^.

Neutrophils are the first line innate immune cell defenders that are recruited from the circulation to the lung alveolar spaces to combat respiratory infection^5^. Unlike much of the periphery where neutrophils emigrate to sites of inflammation through post-capillary venules, in the lung their main site of emigration is the capillary bed^7,8^. Mechanical changes and granule ligand-receptor interactions in the vascular bed contribute to neutrophil retention in the pulmonary capillary network.

Under hemorrhagic conditions, the neutrophils get pre-activated or primed^9^, which in turn obstructs their transit from the intravascular to the extravascular pulmonary spaces^10^. While priming in general is an important event that prepares the immune cells to fully thrust their effector functions against any incoming insult^11^, unregulated priming in conditions like hemorrhage contributes to tissue injury^12,13^. Priming in such conditions leads to release of extracellular proteases and other granule contents in response to an additional stimulus, along with pre-activation of the plasma membrane NADPH oxidase complex. This phenomenon is part of the ‘two-hit hypothesis’ of organ damage^14^ which postulates that neutrophils get primed due to initial trauma or hemorrhage which when followed by a second hit, such as a secondary infection or surgery, leads to premature and overexuberant release of those microbicidal contents. This results in self-injury to the host and leads to acute inflammatory pathogenesis including ARDS^14,15^.

In the current study, we were interested in investigating neutrophil exocytosis during respiratory infection secondary to hemorrhage and how neutrophils could be modulated to improve host outcomes. Because azurophilic granule exocytosis can modulate neutrophil surface properties and inflammatory responses, we investigated whether targeting this process could influence neutrophil trafficking from the pulmonary microcirculation into the extravascular airspaces following hemorrhage and a secondary respiratory insult. In general, the fate of a neutrophil granule, whether it is destined to secrete its cargo in the phagosome or extracellularly at the plasma membrane, depends entirely on its secretory machinery. While delivery of granule contents into the phagosome contributes to intracellular killing of engulfed pathogens^16^, extracellular release of granule proteins can mediate antimicrobial activity in the extracellular milieu and may also contribute to neutrophil extracellular trap (NET) formation and related functions^17,18^. For example, whether an azurophilic granule is associated with plasma membrane exocytosis or phagosomal bacterial killing depends on the presence or absence of the small GTPase Rab27a^19^. Among the various subsets of granules that may be exocytosed by neutrophils, azurophilic granules contain the most toxic bactericidal contents, such as MPO, cathepsin G, elastase and lysozymes. It has been reported that the delivery of azurophilic granule contents into phagosomes concentrates their antimicrobial contents within a confined compartment, promoting efficient intracellular killing of engulfed pathogens^16,20^. It is well established that excessive extracellular release of these bactericidal contents contributes to disease pathogenesis, such as systemic inflammation in sepsis, or acute lung injury^18,21^. Hence, our primary target was to modulate Rab27a such that exocytosis of azurophilic granules, as a hemorrhage-primed event, was prevented while retaining their delivery to the phagolysosomes during the process of neutrophil antimicrobial function at the infection site. We posit that targeting Rab27a could help maintain a balance between the deleterious versus the beneficial effects of the neutrophil response in the context of hemorrhage.

Previous studies have shown that Rab27a-dependent neutrophil exocytosis contributes to inflammatory responses, and that pharmacological disruption of the Rab27a-JFC1 interaction by Nexinhib20 reduces the release of neutrophil granule contents and systemic inflammation in models of endotoxemia^22,23^. Importantly, Nexinhib20 was reported to inhibit azurophilic granule exocytosis without impairing phagocytosis or NET formation in the experimental models examined^24^. Moreover, it was shown that limiting exocytosis during *Streptococcus pneumoniae* lung infection in mice reduced pulmonary barrier injury and bacteremia without affecting the total bacterial load^25^. Based on these findings, our primary target was to modulate Rab27a functions such that it could attenuate hemorrhage-induced neutrophil exocytosis and activation, and excessive inflammation following a secondary respiratory challenge. Given the importance of neutrophil antimicrobial activity in host defense, we also sought to examine the effects of Rab27a on neutrophil-mediated bacterial killing.

To this end, we employed both genetic perturbation and pharmacologic disruption of Rab27a in a mouse model of hemorrhage followed by pulmonary challenge. Our study highlights the potential of therapeutic Rab27a modulation to regulate hemorrhage-induced neutrophil pre-activation and facilitate neutrophil migration from the intravascular compartment to the extravascular spaces of the lung. Consistent with previous findings, we show that Rab27a inhibition attenuates both systemic and pulmonary inflammation under hemorrhagic conditions. In addition, we demonstrate that Rab27a attenuation projects a better bacterial killing capacity under specific *in vitro* conditions. Overall, our findings suggest that Rab27a modulation may help regulate neutrophil dysfunction caused by hemorrhagic shock and potentially benefit the host response to secondary infection.

## Results

### Effect of hemorrhage on neutrophil degranulation and trafficking in the lung

Priming is an important phenomenon that helps neutrophils to prepare their armaments against the onset of any insult to the host^11^. However, hemorrhage-driven partial degranulation of those neutrophils before a microbe encounter serves as a marker of premature activation, beyond neutrophil priming. To understand how prior hemorrhage impacts the neutrophil response to a second insult, mice were subjected to hemorrhage followed 24 h later by *S. aureus* respiratory infection. At 24 h post-infection, whole lungs were excised from one cohort of mice while bronchoalveolar lavage fluid (BALF) was harvested from another cohort (Fig. 1A), to assess neutrophil degranulation (Fig. 1B) and to quantify neutrophil recruitment (Fig. 1C), respectively.

**Figure 1:**
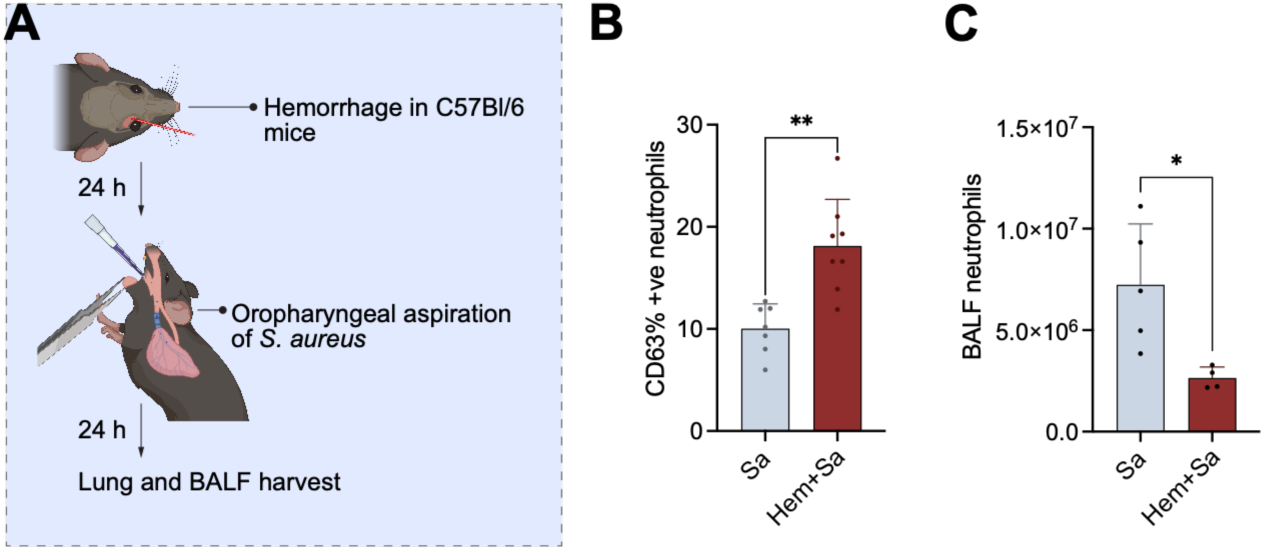
Hemorrhage followed by *S. aureus* infection increases pulmonary neutrophil granule exocytosis and reduces neutrophil availability in the bronchoalveolar lavage fluid. **(A)** Flow diagram of hemorrhage and *S. aureus* lung infection in mice. **(B** and **C)** Following hemorrhage (hem) and secondary respiratory infection in mice using *S. aureus* (Sa), lungs were harvested, homogenized, and analyzed for CD63-expressing Ly6G+ neutrophils by flow cytometry **(B)**, bronchoalveolar lavage fluid (BALF) was harvested from a second set of mice to analyze the total neutrophil (Ly6G+) count by flow cytometry **(C)**. Data are represented as mean±standard error of the mean (SEM). Comparisons between two groups were performed using unpaired two-tailed Student’s t-tests; *p<0.05, **p<0.001

In mice subjected to hemorrhage prior to instillation of *S. aureus* in the lung, the neutrophils from whole lung homogenate exhibited increased expression of CD63, a marker of azurophilic degranulation^19,26^, compared to the infected group with no prior hemorrhage (Fig. 1A, B). Under the same treatment conditions, the hemorrhaged/infected group had decreased neutrophil abundance in the BALF compared to the non-hemorrhaged group (Fig. 1 A, C). While there was no prominent change in weight of mice before and after hemorrhage, there was a significant reduction in their weight post hemorrhage followed by infection (Supplemental Fig. S1). Overall, these results indicate that hemorrhage induces azurophilic degranulation in pulmonary neutrophils and restricts their trafficking into the lung airspaces during an active infection.

### Rab27a deficiency improves neutrophil trafficking from the lung intravascular to the extravascular spaces

We have previously established the methodology for utilizing conditionally immortalized HoxB8 NPs to generate mice that harbor both endogenous and donor-derived neutrophils^27^. NPs are transplanted into mice where they home to the bone marrow, mature into neutrophils, and finally egress to the periphery, and perform effector functions equivalent to host endogenous neutrophils^27^. These NPs can be genetically manipulated to study specific gene functions. Rab27a is a prominent player in maintaining tight regulation of neutrophil vesicular trafficking and azurophilic degranulation^28^. Based on our above observation that azurophilic degranulation is enhanced in a hemorrhage-driven activated neutrophil state, we next sought to target Rab27a and determine whether deficiency of Rab27a could dampen such premature activation and enable better emigration of neutrophils to the lung extravascular spaces from the intravascular regions. To do so, we first knocked out the Rab27a gene from NPs using CRISPR-Cas9 (Supplemental Fig. S2A). After validation of its specific loss-of-function by Western blot (Supplemental Fig. S2B), both wild-type (WT) and the Rab27a knock-out (KO) NPs were transplanted into mice, and 5 days later hemorrhage was inflicted on those mice. This was followed 24 h later by *S. aureus* respiratory infection. To understand the compartment-specific trafficking pattern of the WT versus Rab27a KO neutrophils, the mice were intravenously injected with anti-Ly6G antibody for 5 min, a length of time that is sufficient to stain the intravascular neutrophils without exposing neutrophils outside of the vasculature to the labeling antibody^29^. Following the brief labeling procedure, the circulation was perfused with saline to remove non-sequestered neutrophils and excess antibody, followed by lung harvest, homogenization, and labelling with other antibodies for analysis by flow cytometry (Fig. 2A). Flow cytometry analyses were performed to distinguish intravascular from extravascular neutrophils and to identify donor-derived neutrophils (Supplemental Fig. S3). The fraction of KO neutrophils out of all donor neutrophils was higher in the extravascular space compared to the intravascular regions as opposed to WT neutrophils which were seen to be more abundant in the pulmonary capillaries and less available in the extravascular space compared to Rab27a KO (Fig. 2B). Although not as pronounced, this trend was also seen in mice that were not subjected to hemorrhage prior to infection, suggesting that Rab27a KO neutrophils may have better emigration ability in the infected lungs generally (Fig. 2C).

**Figure 2:**
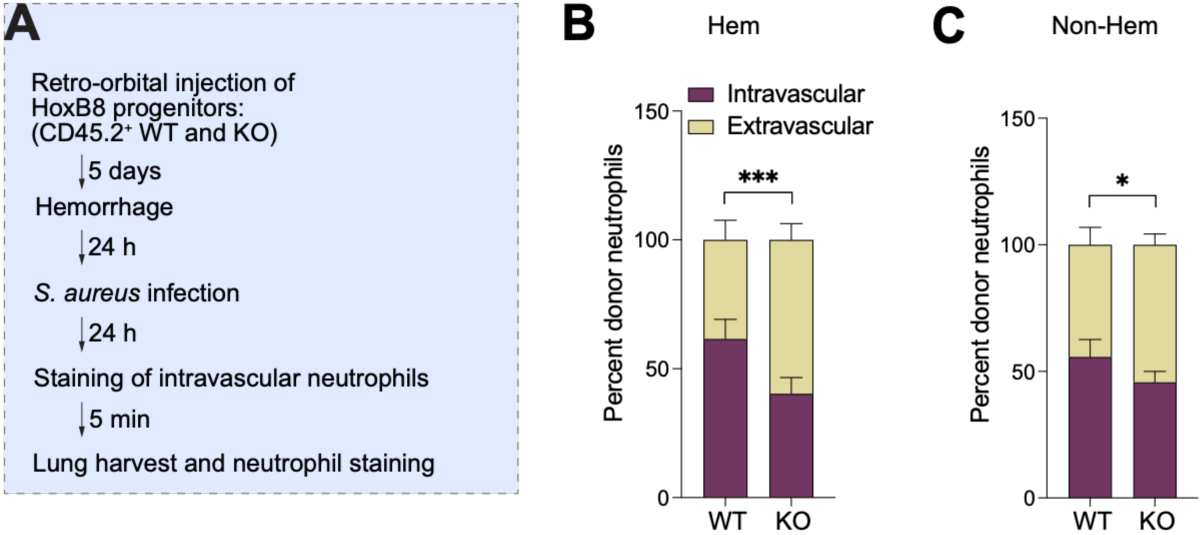
Rab27a deficiency improves trafficking of neutrophils into the lung extravascular spaces. **(A)** Flowchart of mice transplantation, treatment, harvest, and staining to understand neutrophil trafficking in lungs by flow cytometry. **(B-C)** Comparison of WT and KO donor neutrophils percentage in intravascular vs extravascular lung regions in hemorrhaged (Hem) **(B)** and non-hemorrhaged (Non-Hem) **(C)** mice infected with *S. aureus*. Data are represented as mean±SEM. Two-way analysis of variance (ANOVA) was performed with Tukey’s multiple comparison test; each group n=5 mice, *p<0.05, ***p<0.0001

### Effect of Nexinhib20, a small molecule inhibitor of Rab27a on neutrophil trafficking to lung airspaces

Enhanced trafficking of Rab27a-deficient neutrophils into the lung extravascular spaces of hemorrhaged mice suggests a potential therapeutic strategy involving modulation of Rab27a-associated functions that may help overcome impaired neutrophil availability at the site of an active respiratory insult, i.e. the lung airspaces of those mice. Consequently, we next investigated the effects of pharmacological inhibition using Nexinhib20. Nexinhib20 is a small molecule inhibitor that hinders Rab27a-mediated neutrophil granule exocytosis by preventing the binding of Rab27a with its effector protein, JFC1^24^. Nexinhib20 was reported as an inhibitor of azurophilic granule exocytosis and extracellular superoxide anion production in human neutrophils, without affecting phagocytosis or NETosis^24^. Hence, use of this small molecule inhibitor would enable the acute modulation of Rab27a associated exocytosis without the need for genetic manipulation. It was also shown that Nexinhib20 reduced systemic inflammation in mice in response to intraperitoneal injection of LPS^24^. Hence, in our next experiment, we used LPS as a defined inflammatory stimulus to assess effects of Nexinhib20 on host inflammatory responses eliminating the variable of bacterial burden that could be impacted by Nexinhib20. To do so, mice received intraperitoneal injection of Nexinhib20 before undergoing hemorrhage, followed 24 h later by respiratory instillation of LPS. The mice were evaluated after 3 h of LPS challenge (Fig. 3A). As expected, we observed reduced neutrophil numbers in the BALF of the hemorrhage-LPS group compared to the sham-LPS mice, and although the values could not reach significance Nexinhib20 treatment was able to partially restore neutrophil recruitment under hemorrhagic conditions (Fig. 3B). In addition, we assessed neutrophil exocytosis in these conditions by analyzing the surface expression of CD63, and CD35, a complement receptor and a marker for secretory vesicles^24,30^, both in the BALF and blood using flow cytometry. The BALF neutrophils showed a potentially increasing trend of CD63 surface expression in the hemorrhage-LPS group compared to the sham-LPS mice, although the values could not reach significance. Further, Nexinhib20 significantly reduced the fraction of those CD63 positive neutrophils induced by hemorrhage-LPS (Fig. 3C). A similar kind of trend was also seen in the blood neutrophils (Supplemental Fig. S4A) although the values did not reach significance. However, CD35 surface expression was barely detectable in both the blood and BALF neutrophils at the given time-point (with MFI ranging from 0.5 to 2.5, Supplemental Fig. S4B, C). Overall, these observations indicate that Nexinhib20 mimics Rab27a KO phenotype by potentially improving neutrophil availability, and by reducing neutrophil azurophilic exocytosis in a hemorrhage-LPS two-hit mouse model.

**Figure 3:**
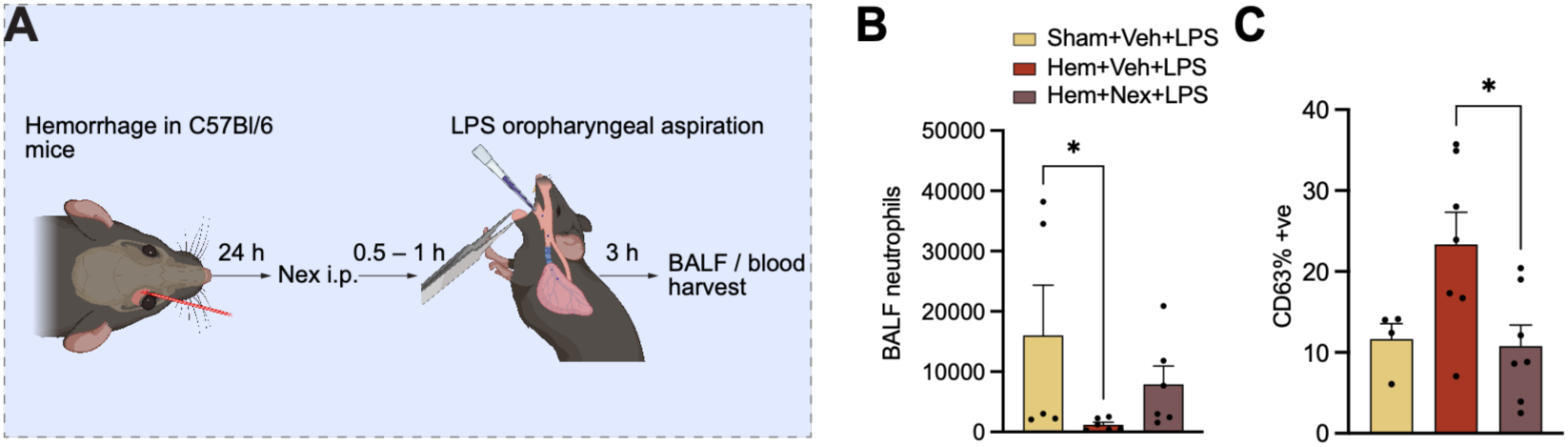
Effect of LPS challenge and Nexinhib20 treatment on the phenotype of bronchoalveolar lavage neutrophils following hemorrhage. **(A)** Flow diagram of Nexinhib20 (Nex) treatment of mice subjected to hemorrhage (hem) and LPS oropharyngeal aspiration. **(B–C)** After hemorrhage followed by Nexinhib20 (Nex) or vehicle (Veh) treatment, and LPS instillation for the given time-point, BALF was harvested to analyze the total neutrophil (Ly6G+) count **(B)** and to assess neutrophils expressing surface CD63 by flow cytometry **(C)**. Data are represented as mean±SEM. Data normality was assessed using the Shapiro-Wilk normality test. Non-parametric Kruskall Walis test or ordinary one-way ANOVA was done with Dunnett’s multiple comparison test; *p<0.05

### Effect of Rab27a modulation on systemic and pulmonary inflammation

Although neutrophils are the first line defender immune cells to resolve any kind of infection, increased neutrophil infiltration contributes to inflammation in several pathogenesis. For instance, higher neutrophil influx in host tissues of patients post-trauma is considered an indicator of heightened inflammation^31^. Enhanced neutrophil infiltration has also been reported in models of systemic inflammation such as colitis or diabetes causing damage to the tissues^32^. However, in the present context where neutrophil influx into the airspaces, the primary site of respiratory insult, is restricted, maintaining sufficient neutrophil availability while limiting excessive inflammation would be beneficial to the host. To this end, we used blood plasma and BALF samples collected from the previous experiment (Fig. 3A) to evaluate important markers associated with host inflammation, and whether Nexinhib20 could attenuate systemic and pulmonary inflammatory responses under these conditions. Among these, MPO, a major enzyme stored in neutrophil azurophilic granules^18^, was of particular interest because circulating MPO levels have been associated with endothelial dysfunction, and as a potential biomarker of disease severity and sepsis^32,33^. Although MPO derived oxidants including hypochlorous acid (HOCl) contribute to innate antimicrobial defense, excessive MPO release and its oxidative products may promote vascular and tissue damage, such as in atherosclerotic lesion development^33^. Furthermore, previous studies have reported that Nexinhib20 reduces circulating MPO levels in a model of systemic inflammation induced by intraperitoneal LPS administration^24^. Based on these findings, we evaluated plasma MPO concentrations by ELISA in our model of hemorrhage followed by respiratory LPS challenge. As expected, there was higher circulating MPO in the hemorrhage-LPS group compared with naive controls. Although the hemorrhage-LPS group showed an increasing trend also when compared to the LPS group that received sham procedure, the values were not significant. Notably, Nexinhib20 treatment significantly reduced the circulating MPO levels compared with the untreated hemorrhage-LPS group (Fig. 4A). Thus, pharmacological intervention using Nexinhib20 attenuated the systemic MPO response in this model.

**Figure 4:**
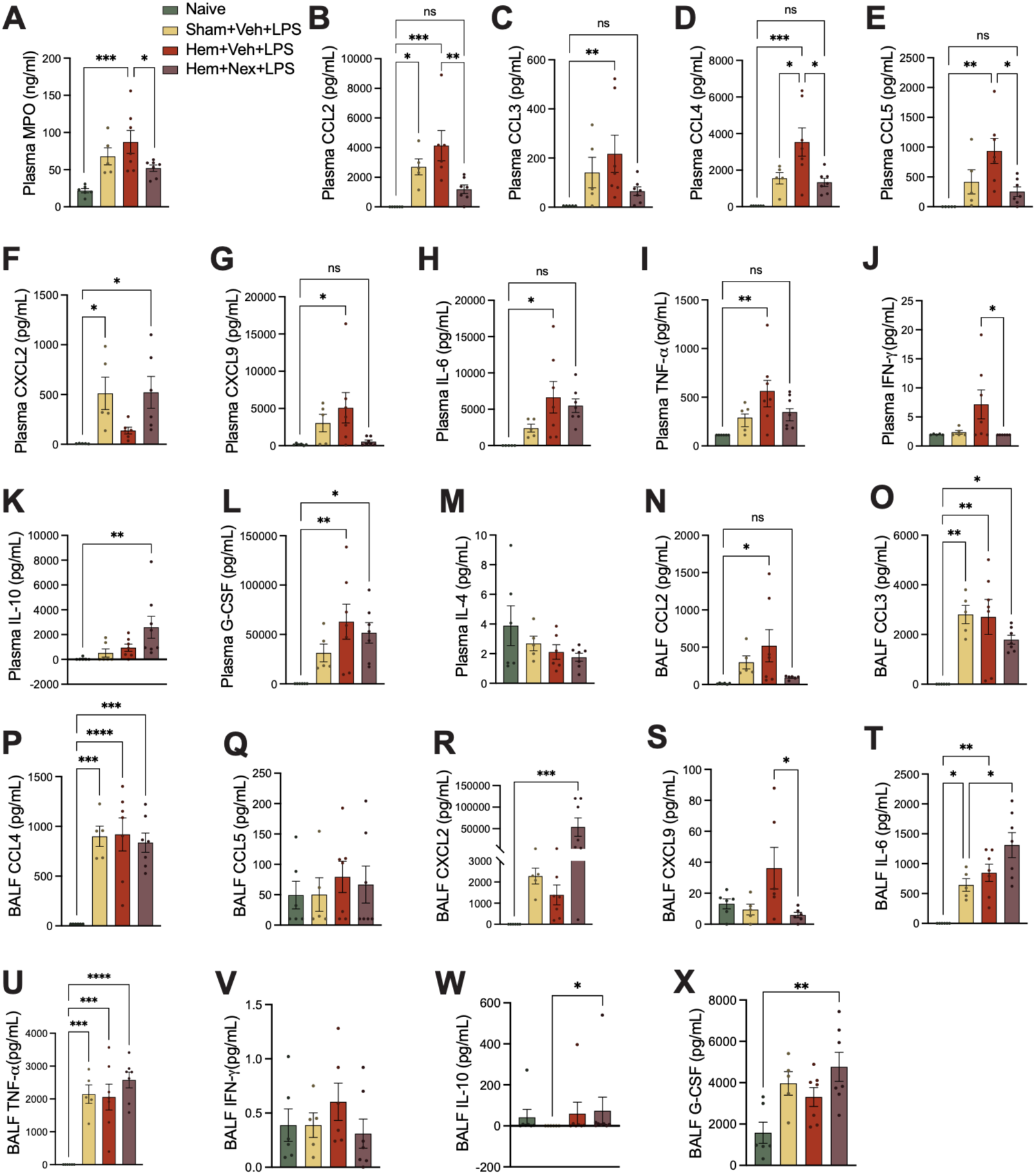
Nexinhib20 differentially modulates systemic and pulmonary inflammation following hemorrhage and oropharyngeal LPS challenge. **Mice were** subjected to hemorrhage (hem) followed by Nexinhib20 (Nex) or vehicle (Veh) treatment and LPS oropharyngeal aspiration after 24 h. Mice were sacrificed after 3 h of LPS administration, and blood plasma, BALF were harvested to analyze plasma myeloperoxidase (MPO) levels by ELISA **(A)**, inflammatory mediator levels in blood plasma **(B-M)**, and BALF **(N-X**) by LEGENDplex^TM^ multiplex immunoassay (BioLegend). Data are represented as mean±SEM. Data normality was assessed using Shapiro-Wilk normality test. One-way ANOVA was performed with Dunnett’s or Tukey’s multiple comparison test. When assumptions of normality were not met, the nonparametric Kruskal–Wallis test followed by Dunn’s multiple-comparisons test was used for one-way analyses. *p<0.05, **p<0.01, ***p<0.001, ****p<0.0001, ns=non-significant (p>0.05).

Next, we analyzed levels of inflammatory mediators in the blood plasma (Fig. 4B-M) and BALF (Fig. 4N-X). In the blood plasma, levels of chemokines CCL2, CCL3, CCL4, CCL5, CXCL9, as well as cytokines IL-6, TNF-α, were significantly elevated in the hemorrhage-LPS group compared to naïve control (Fig. 4B-E, G-I). Out of these, CCL2, CCL4, CCL5 were significantly reduced by Nexinhib20 treatment. CCL3 and CXCL9 levels trended towards potentially lower values with Nexinhib20 treatment but could not reach significance (Fig. 4B-E, G). Although IL-6 and TNF-α levels in the Nexinhib20 treated hemorrhage-LPS group were not significantly different from those in naive mice, their difference with the untreated hemorrhage-LPS group did not reach statistical significance (Fig. 4H, I). In addition, compared to naive, only CCL2 levels were significantly elevated by LPS insult alone (Fig. 4B). The rest of the mentioned mediators also showed an increasing trend in the sham-LPS group although the levels did not reach significant values. Furthermore, while all of these, showed an increasing trend in the hemorrhage-LPS group compared to their sham-LPS counterparts, only CCL4 showed a significant increase (Fig. 4B-E, G-I). Interestingly, a key neutrophil migration player, CXCL2^34^ was significantly elevated in the sham-LPS group but was not increased in the plasma of mice subjected to hemorrhage prior to LPS administration. Notably, Nexinhib20 treatment of the hemorrhage-LPS group showed significantly elevated levels compared to naive mice, and the concentrations were comparable to the sham-LPS group (Fig. 4F). In addition, plasma IFN-γ levels showed an increasing trend in the hemorrhage-LPS group only, and that was significantly brought down by Nexinhib20 treatment (Fig. 4J). Furthermore, plasma IL-10 was significantly increased by Nexinhib20 compared to the naive group (Fig. 4K). However, IL-4 levels were near the limit of detection with no significant changes across the mice groups (Fig. 4M). In addition to these chemokines and cytokines, G-CSF also showed a significantly elevated level in the hemorrhage-LPS group but not in the sham-LPS group compared to naive mice, however, Nexinhib20 did not exert a significant reduction of those increased levels (Fig. 4L).

Under the same conditions, inflammation profile analysis of the BALF showed significant increase of CCL2, CCL3, CCL4, IL-6, and TNF-α in the hemorrhage-LPS group compared with naive mice. Out of these, in the Nexinhib20 treated hemorrhage-LPS group, only CCL2 levels were no longer significantly different from those in naive mice, although the difference between the treated and untreated hemorrhage-LPS groups did not reach statistical significance (Fig 4N-P, T, U). Furthermore, contrary to significant CCL2 elevation by LPS alone in plasma (Fig. 4B), the BALF CCL2 levels were not significantly raised in the sham-LPS group (Fig. 4N). In addition, contrary to changes observed in plasma CCL5 levels (described above), BALF CCL5 levels remained comparable across all experimental groups (Fig. 4E, Q).

Interestingly, compared to naive, CXCL2 showed a potentially increasing trend in both LPS groups, with and without hemorrhage, although the values did not reach significance, and was significantly elevated only in the Nexinhib20 treated hemorrhage-LPS group (Fig. 4R). Next, CXCL9 levels showed a potentially increasing trend only in the hemorrhage-LPS group compared to naive, although the values did not reach significance, and those levels were significantly reduced upon Nexinhib20 treatment (Fig. 4S). Further, negligible levels of IFN-γ were detected at the given time-point with no change across the groups (Fig. 4V). The overall levels of IL-10 were also very low or undetected in the BALF, with the Nexinhib20 treated group exhibiting an increase compared to the sham-LPS group (Fig. 4W). In addition, IL-4 levels were undetected in the BALF in the given time and treatment conditions (and therefore, data not shown). Although there was an increasing trend of BALF G-CSF in all the other groups compared to naive, significance of this pattern was only observed in the Nexinhib20 treated hemorrhage-LPS group (Fig. 4X). Collectively, these findings demonstrate that hemorrhage followed by an acute respiratory challenge induces systemic and pulmonary inflammatory responses as early as 3 h. Furthermore, pharmacological treatment with Nexinhib20 attenuated several of these inflammatory responses, highlighting the potential of targeting Rab27a to modulate inflammation under such conditions.

### Nexinhib20-mediated Rab27a modulation and the bacterial killing capacity of neutrophils

Regulating detrimental effects of inflammation without hindering the ability of neutrophils to eradicate microbes is a desirable outcome for host defense. In previous studies, Nexinhib20 was reported to inhibit neutrophil exocytosis without affecting other innate immune effector functions, such as phagocytosis and NETosis^24^. However, to our knowledge, the effects of Nexinhib20 on neutrophil-mediated bacterial killing during infection have not been directly investigated. To understand this aspect, we employed an *in vitro* killing assay using neutrophils and bi-fluorescent *S. aureus*. This assay involves the use of *S. aureus* stably expressing green fluorescent protein (GFP) that is further labelled with a secondary fluorescent dye conjugate, ATTO647, a highly stable molecule whose fluorescence is not impacted even after the bacterium has been internalized and killed within the phagolysosome. This allows us to differentiate the phagocytosing neutrophils containing live GFP expressing ATTO positive bacteria that have not yet been killed, from the neutrophils that are GFP negative ATTO positive indicating the population that has completed killing of the internalized bacteria^35^ (Fig. 5A, B). First, to validate the inhibitory effect of Nexinhib20 on granule exocytosis, we measured the surface expression of CD63 in all Ly6G+ATTO+ neutrophils at 90 min after exposure to bi-fluorescent *S. aureus*, the time-point at which the killing efficacy was assessed in subsequent experiments. The Ly6G+ATTO+ population represents all phagocytosing neutrophils^35^. As expected, the cell surface CD63 expression was suppressed in the Nexinhib20 treated group compared to the untreated Ly6G+ATTO+ neutrophils (Fig. 5C). Next, we assessed the percentage of ATTO+GFP-cells in the Ly6G+ population which represents the number of neutrophils that have phagocytosed and completed the killing of *S. aureus*. We found that the fraction of ATTO+GFP-neutrophils was higher in the Nexinhib20 treated group compared to the control group at 90 min (Fig. 5D). We then calculated the killing index as the fraction of neutrophils that had completed killing of internalized *S. aureus* among all neutrophils that were ATTO+, as described elsewhere^35^. The killing index of the Nexinhib20 treated group was significantly higher than the untreated group at 90 min (Fig. 5E). Together, these data indicate that Nexinhib20 enhances the intracellular eradication of *S. aureus* by neutrophils. In addition, we went on to check whether Nexinhib20 affects the overall extent of *S. aureus* phagocytosis, by measuring the ATTO MFI, and although there was an increase in phagocytosis level from 0 to 90 min in the control group, there was no significant increase in ATTO MFI for the Nexinhib20 treated group at 90 min compared to 0 min (Fig. 5F). This suggests that Nexinhib20 may limit the extent of phagocytosis, which could contribute to the ability of neutrophils to contain and kill the bacteria that they have already engulfed. However, the fraction of ATTO+ neutrophils did not change across the groups over time (Fig. 5G), indicating that the overall number of neutrophils participating in phagocytosis is not impacted by Nexinhib20. Taken together, while Nexinhib20 prevents extracellular degranulation of azurophilic granules, it may improve the intracellular bacterial killing capacity of those neutrophils.

**Figure 5:**
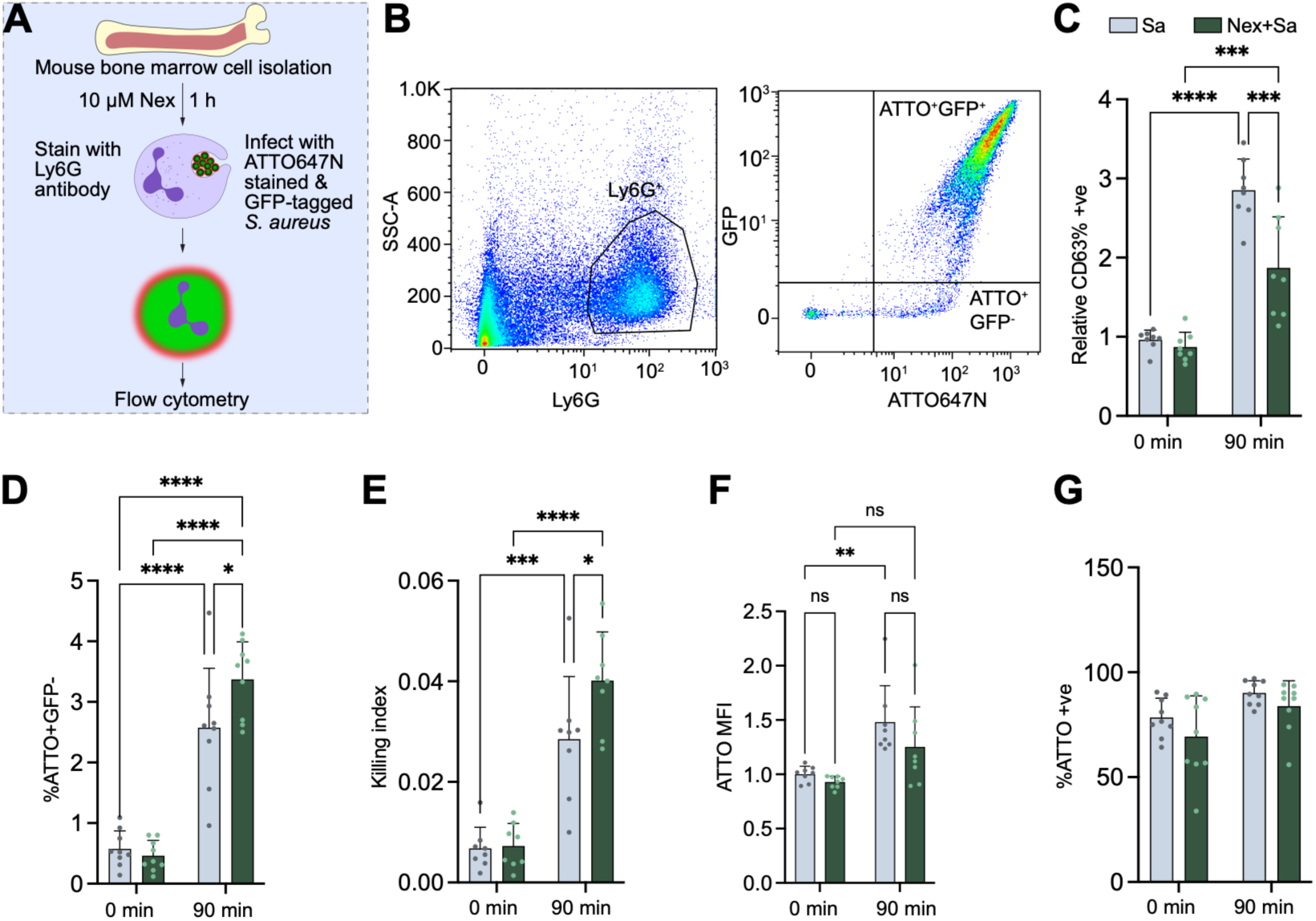
Nexinhib20 enhances neutrophil-mediated *S. aureus* killing under specific *in vitro* conditions. **(A)** Schematic of *in vitro* treatment of bone marrow cells with Nexinhib20 (Nex) and *S. aureus*. **(B–G)** Bone marrow cells were pre-treated with Nex (10 μM) for 1 h followed by infection with GFP-tagged *S. aureus* (Sa) (MOI 10) dyed with ATTO647N for 0 min and 90 min. Cells were stained with anti-Ly6G and anti-CD63 antibodies and analyzed by flow cytometry. Two-way ANOVA was performed with Tukey’s multiple comparison test; *p<0.05, **p<0.01, ***p<0.001, ****p<0.0001, ns=non-significant (p>0.05).

### Nexinhib20 induces retention of neutrophil intracellular MPO

In general, granules fuse with the phagosomal membrane to deliver catalytic or non-catalytic antimicrobial cargoes for phagocytosis^36^. Specifically, the azurophilic granule protein MPO^18^ catalyzes the production of hypochlorous acid (HOCl) that is highly toxic to the bacteria, from H_2_O_2_ and chloride, inside the phagosome^37^. Schwartz et al. and others have shown that exogenously expressed cytosolic GFP by bacteria is bleached by this MPO-generated HOCl in neutrophils during an infection^36,38^. To understand the potential mechanisms by which Nexinhib20 enhances bacterial killing in our study using bi-fluorescent *S. aureus* (Fig. 5), we employed flow cytometry to detect intracellular MPO levels under similar treatment conditions. Briefly, bone marrow cells were treated with Nexinhib20, stained with Ly6G antibody, followed by exposure to GFP-expressing *S. aureus* for 90 min. The cells then underwent intracellular staining with MPO and were analyzed by flow cytometry. As expected, GFP signal was reduced in the Ly6G+ neutrophils of the Nexinhib20 treated group at the 90 min time-point compared to the control group (Fig. 6A), indicating enhanced bacterial killing. Interestingly, the MPO levels remained constant over the time period of 0 min to 90 min in the Nexinhib20 treated group, as opposed to the control group where the intracellular MPO levels significantly decreased from 0 to 90 min (Fig. 6B, C). Accordingly, the percentage of MPO-high neutrophils remained constant from 0 to 90 min in the Nexinhib20 group while the analogous population decreased in frequency over time in the control group (Fig. 6D). Together, these findings suggest that Nexinhib20 treatment promotes neutrophil intracellular MPO retention during *S. aureus* infection, potentially by limiting extracellular MPO release from those neutrophils.

**Figure 6:**
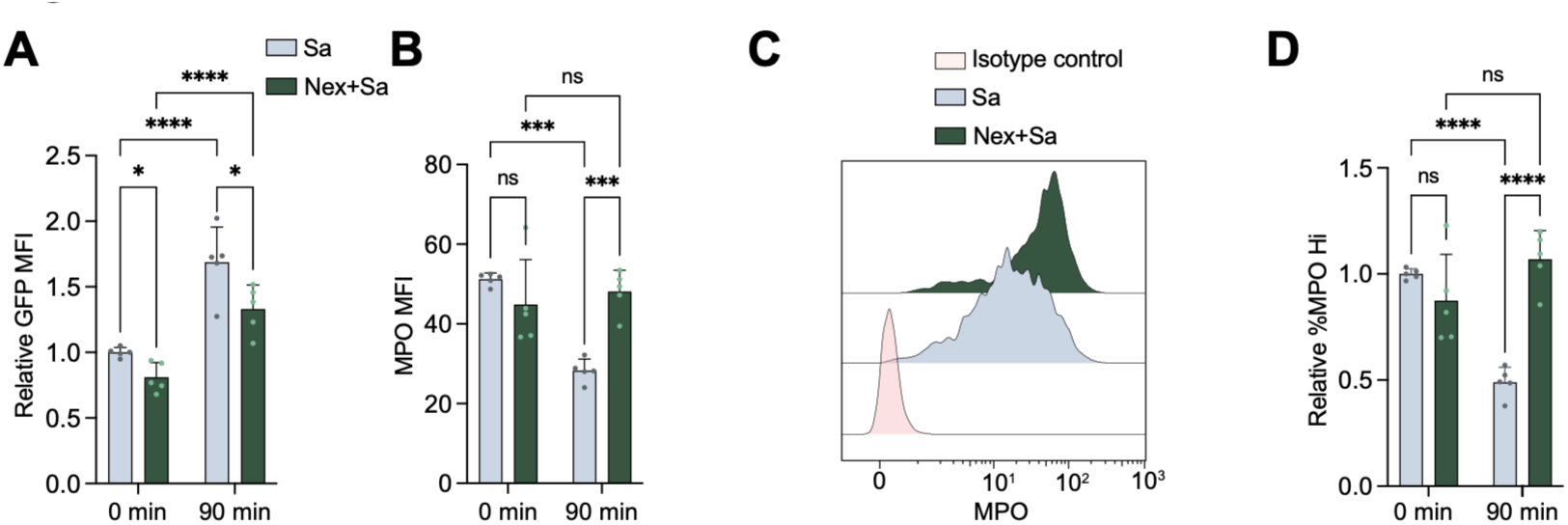
Nexinhib20 promotes intracellular MPO retention in neutrophils during *S. aureus* infection *in vitro*. Bone marrow cells were pre-treated with Nexinhib20 (Nex) (10 μM) for 1 hour followed by infection with GFP-tagged *S. aureus* (Sa) (MOI 10) for 0 min and 90 min. Cells were surface-stained with anti-Ly6G followed by intracellular staining with anti-MPO or isotype control for analysis using flow cytometry. Two-way ANOVA was performed with Tukey’s multiple comparison test; *p<0.05, ***p<0.001, ****p<0.0001, ns=non-significant (p>0.05).

## Discussion

Susceptibility to secondary infections is a common manifestation of the immune dysfunction in critically ill patients following trauma-related hemorrhage^39,40^. Methicillin-resistant *S. aureus* contributes to 20-40% of all nosocomial pneumonias in such patients^41^. While neutrophils are the first line defender immune cells during an acute bacterial infection, their availability in the airspaces can be restricted by widespread premature activation or priming as a result of hemorrhage^7,42–44^. Although priming, in general, is a prerequisite for immune cells to prepare their armaments against any insult^45,46^, unwanted or untimely priming could do more harm than good to the host^46–48^. Using a mouse model of hemorrhage and secondary respiratory infection with *S. aureus*, we found that neutrophils prematurely degranulate (exocytose prepackaged granules) while still in the intravascular capillary network of the lungs. Owing to such premature activation in the pulmonary microcirculation, the neutrophils emigrate less readily into the airspaces, the active site of infection. Consistent with the observations here, we have previously shown that prior hemorrhagic shock results in attenuated neutrophil recruitment into the airspaces in response to both *Pseudomonas aeruginosa*^49^ and *S. aureus*^50^. At the outset, one goal of this study was to identify mechanisms underlying the changes in neutrophil trafficking in the lungs that occur downstream of hemorrhage. Using genetic disruption of Rab27a and its pharmacologic inhibition with Nexinhib20, we have found here that hemorrhage-primed release of azurophilic granules contributes to restricting neutrophil entry into the infected airspaces of the lung.

The small GTPase Rab27a is a key determinant of neutrophil extracellular degranulation or exocytosis of azurophilic granules, the granule subset containing the most toxic bactericidal cargoes^19^. Using neutrophils from Rab27a KO mice it was shown that MPO secretion from azurophilic granules was completely impaired in those neutrophils. In humans, defects in this gene lead to a rare genetic disorder called Griscelli syndrome type 2 (GS2) that occurs due to defects in melanasome transport and causing impaired functioning of cytotoxic T cells, neutrophils, and natural killer cells. In absence of Rab27a, the azurophilic granules remain able to be directed towards the phagosome for intracellular bacterial killing in response to appropriate stimuli^51,52^. In addition, it is known that excessive neutrophil exocytosis in the bloodstream contributes to the pathophysiology of sepsis^53–55^. Hence, our primary goal was to inhibit exocytosis while retaining intracellular degranulation and other anti-pathogen effector functions of the neutrophils. To this end, we found that knockout of Rab27a promoted neutrophil transit from the intravascular lung regions to the extravascular spaces in the *S. aureus* infected mice, both in the absence or presence of prior hemorrhage. These observations indicate that lack of Rab27a associated pre-activation of the neutrophils helps in better regulation of their activation status during an insult.

Further, we took advantage of the availability of a small molecule inhibitor of Rab27a, called Nexinhib20 to study its effects on mitigating inflammatory symptoms in the host under such two-hit insult conditions. To specifically investigate host defense mechanisms independent of the confounding effects of bacterial viability and replication, we employed LPS as a defined bacterial-derived inflammatory stimulus rather than whole live bacteria. We found that hemorrhage significantly reduced BALF neutrophil numbers following respiratory LPS challenge compared with the sham-LPS group, suggesting impaired neutrophil accumulation in the pulmonary airspaces under hemorrhage-LPS dual insult. Pharmacological inhibition of Rab27a using Nexinhib20 resulted in a numerical increase of the BALF neutrophils compared with the untreated hemorrhage-LPS group, although the difference did not reach statistical significance. This trend was directionally consistent with our Rab27a KO phenotype that showed enhanced pulmonary extravascular trafficking of those KO neutrophils. This observation fortified our understanding that irrespective of the insult type and the type of Rab27a perturbation, either in the form of genetic depletion in our donor neutrophil progenitors (as in Fig2), or small molecule inhibition by Nexinhib20 (Fig3), attenuation of Rab27a could help in better trafficking of the neutrophils to the site of active insult. In addition, consistent with previous reports, Nexinhib20 reduced systemic plasma MPO levels in the hemorrhage-LPS mice. Furthermore, Nexinhib20 administration attenuated several systemic and pulmonary inflammatory mediator responses. Particularly, hemorrhage/LPS-induced circulating plasma levels of CCL2, CCL4, CCL5 were significantly reduced by Nexinhib20. Moreover, in the BALF, only CXCL9 was significantly reduced in the Nexinhib20 treated hemorrhage-LPS mice group, while CCL2 showed a potentially reducing trend. Interestingly, higher CXCL2 concentration levels as observed in the BALF compared to that in the plasma, and more so in the BALF of Nexinhib20 treated hemorrhage-LPS group, is indicative of a potential concentration gradient to attract more neutrophils to the airspaces. In previous studies, CXCL2 has been reported to drive neutrophil recruitment to the lung during an infection, for instance in COVID-19 patients^56^, or in acute lung injury^57^. Furthermore, Nexinhib20 treatment resulted in a favorable increase of the anti-inflammatory IL-10 as observed both in the plasma and the BALF of mice subjected to hemorrhage and LPS exposure, indicative of enhanced anti-inflammatory signaling in response to the given experimental conditions. Overall, these observations indicate that respiratory challenge using LPS triggers both systemic and pulmonary inflammation as early as 3 h, and curbing of Rab27a-driven neutrophil exocytosis using pharmacological intervention attenuates the host inflammatory response. In addition, while most of the pro-inflammatory readouts were attenuated by Nexinhib20, the neutrophil recruitment into the airspaces was supposedly improved in the BALF. This suggested that, in contrast to the conventional interpretation of increased neutrophil numbers as a marker of inflammation^58,59^, neutrophil trafficking is also an important determinant of their localization and potential functions. In the setting of hemorrhage followed by a secondary respiratory insult, the release of neutrophils sequestered in the pulmonary microcirculation and their subsequent migration into the airspaces may be critical for their access to the site of active insult.

It has been previously established that Nexinhib20 inhibits Rab27a-regulated exocytosis without affecting other innate immune functions of neutrophils, such as phagocytosis or NETosis^24^. It is also reported to inhibit upregulation of adhesion molecules such as CD11b^24^, however, in our experimental setup Nexinhib20 had no significant effect on CD11b surface expression on neutrophils in the hemorrhage-LPS group (Supplemental Fig. S5). This could be due to the fact that while CD11b serves as a standard marker for mature mouse neutrophils alongside Ly6G^60^, its surface expression is also used to designate specific granules and secretory vesicles in stimulated neutrophils^61^. Due to these, the occurrence of expression changes of CD11b could be masked in our given experimental conditions. As previously described, CD35, a marker for neutrophil secretory vesicles, was also barely detectable on circulating and BALF neutrophils at the assessed time-point (Supplemental Fig. S4B-F). These may reflect differences in the timing or extent of granule mobilization under the experimental conditions examined.

In addition, using bone marrow neutrophils we show that while Nexinhib20 reduced surface CD63, it augmented, under some conditions, the killing efficacy of the phagocytosed *S. aureus*. Delving deeper into its underlying mechanism, we found that Nexinhib20 retains the intracellular MPO levels in this condition, whereas control neutrophils exhibited a decrease in intracellular MPO when stimulated with *S. aureus*. In the phagosome, MPO is critical for generating species toxic to microbes, including HOCl^62,63^. Previous reports have shown that MPO-generated HOCl bleaches GFP tagged to the phagocytosed *S. aureus* in human neutrophils^36^. Thus, our observations suggest that the retention of MPO in Nexinhib20 treated neutrophils may mechanistically contribute to enhanced *S. aureus* killing by the neutrophils. Using the bi-fluorescent *S. aureus* assay, our observation that Nexinhib20 does not significantly impact the fraction of ATTO+ neutrophils coincides with previous reports that Nexinhib20 treatment does not affect neutrophil phagocytosis^24^.

Taken together, the present study shows that Rab27a modulation helps in better regulation of granule exocytosis for effective delivery of neutrophils to the site of active insult in a two-hit mouse model involving hemorrhage and a secondary respiratory stimulus. In addition, we show that pharmacological intervention using Nexinhib20 attenuates the systemic and pulmonary inflammation. Furthermore, under certain conditions, Nexinhib20 also improves *S. aureus* killing efficacy of the neutrophils. Hence, targeting Rab27a may offer a potential strategy to regulate excessive inflammation while preserving or enhancing antimicrobial activity. Further studies are warranted to establish the underlying mechanisms and determine whether these effects translate into improved host defense *in vivo*.

## Materials and methods

### Animals

C57BL/6 male mice 8-12 weeks old were used for all experiments (The Jackson Laboratory, Bar Harbor, ME). All the experiments were conducted with approval from the Brown University Health Institutional Animal Care and Use Committee (Office of Laboratory Animal Welfare Assurance #A3922-01) in accordance with Public Health Service guidelines for animal care and use. Water and standard chow were available ad libitum.

### Fixed-volume hemorrhage model

Mice were anesthetized with isoflurane and blood was withdrawn through the retro-orbital sinus at 40% volume calculated as a percentage of the total blood volume^64^. The total blood volume of a mouse is approximately 69±7 mL/kg of body weight^65^. After hemorrhage, mice were monitored for return to normal activity. A single dose of buprenorphine (0.5 mg/kg) was subcutaneously administered to all mice in the study to alleviate potential pain.

### Nexinhib20 treatment

Mice with or without hemorrhage were first i.p. injected with Nexinhib20 (30 mg/kg) or vehicle. Nexinhib20 was dissolved in DMSO to prepare 100 mM stock concentrations, and was mixed with the co-solvent, 10% Trappsol (2-hydroxypropyl-β-cyclodextrin) (from CTD, Inc., USA) in PBS for *in vivo* administrations.

### Acute respiratory challenge

Methicillin resistant *S. aureus* (strain USA300) was cultured to the log phase in a tryptic soy broth medium. At 0.8 OD_600_, the bacterial culture was pelleted by centrifugation at 3220xg for 10 min, followed by two washes with PBS. Based on 0.5 OD600 being equivalent to 2×10^8 CFU/mL, mice were infected with 75×10^6 CFU in a 50 μL volume. For experiments using LPS, 7.5 mg/mL of LPS in 50 μl PBS was administered to the mice. Mice were anesthetized briefly by isoflurane inhalation and then placed on an intubation stand (Braintree Scientific). The bolus was pipetted at the back of the oral cavity while gently pinching the nose to prevent air passage through the nasal cavity, allowing the bolus to be inhaled via the trachea. After instillation, the mice were immediately allowed to resume normal breathing and returned to their respective cages.

### BALF and blood plasma harvest

BALF was collected postmortem by pushing 1 mL of PBS containing 5 mM EDTA and 0.5% BSA (only PBS was used to harvest BALF for protein content studies) gently into an angiocatheter inserted into the trachea and fixed to a 1 mL syringe, and then gently but steadily pulling fluid back into the syringe. This process was repeated 2-3 times, and the collected BALF was stored in ice until processed for next steps. For ELISA and cytokine assays, the BALF was aliquoted and immediately stored at −80°C for future use.

To isolate plasma, cardiac blood was collected in EDTA microvettes, spun down at 2000xg for 15 min, and the supernatant was immediately isolated and stored in aliquots at −80°C.

### Cytokine bead array

BALF and blood plasma were harvested as described above. The BALF was spun down at 400xg for 10 min to remove cells and debris. Both BALF and plasma were stored in aliquots at −80°C until further use. Cytokine bead array was performed against selected targets using the BioLegend LegendPlex^TM^ as per manufacturer’s protocol.

### NP cell culture and transplantation

Generation of HoxB8-dependent murine NPs has been described previously^27^. Lentivirus was used to transduce NPs to express Cas9 and BFP2 fluorescence. LentiCas9-Blast was a gift from Feng Zhang (Addgene plasmid #52962), and lentiGuide-Hygro-mTagBFP2 was a gift from Kristen Brennand (Addgene plasmid #99374). Lentiviral plasmid containing sgRNA against Rab27a and mCherry was purchased from Genecopoeia (Rockville, MD, USA). After lentiviral transduction, NPs were subjected to single-cell sorting based on double positivity for mCherry and BFP2 into a 96-well plate using BDFACSAria. Rab27a knockout NP clones were identified by Sanger sequencing and validated by Western blotting.

All NPs were maintained in culture using OptiMEM medium (Gibco) supplemented with 10% FBS (Gemini Bio), 1X penicillin-streptomycin (Gibco), 1X non-essential amino acids (Gibco), 30 μM beta mercaptoethanol (Sigma-Aldrich), Z-4 hydroxytamoxifen (4-OHT) (100nM) (Tocris) and stem cell factor (SCF) (50 ng/mL). SCF was obtained from the supernatant of CHO cells (that secrete recombinant murine SCF), a kind gift from Patrice Dubreuil, Centre de Recherche en Cancérologie de Marseille.

Prior to transplantation, the NPs were washed 3-4 times with 1X PBS to remove 4-OHT, and then resuspended to obtain 25-40 million cells in 100 μL PBS. This suspension was transplanted into each mouse through the retro-orbital sinus under isoflurane anesthesia.

### NP cell differentiation

For differentiation of NPs, the cells were first thoroughly washed 3-4 times with 1X PBS to remove 4-OHT, followed by differentiation in media supplemented with SCF (20 ng/mL) and G-CSF (20 ng/mL) (BioLegend) for 2 days, followed by 2-4 days of culture in media supplemented only with G-CSF.

### Staining for intravascular neutrophils

For experiments analyzing neutrophil emigration from intravascular to extravascular lung spaces, WT NPs were stained prior to transplantation with 5 μM CFSE for 20 min at 37°C. The staining was quenched by adding media containing 10% FBS followed by extensive washing with PBS. A 50:50 mixture of WT and KO (unstained) NPs was retro-orbitally injected into the mice under isoflurane anesthesia. On day 5 post-transplant, the mice were subjected to hemorrhage, followed by *S. aureus* oropharyngeal aspiration on day 6. Finally, the mice were sacrificed after 24 h of infection. Right before sacrifice, mice were anesthetized with ketamine/xylazine (100-150 mg/kg, 10 mg/kg), then intravenously injected with APC/Fire750 anti-Ly6G (1 μg in a 100 μL volume). After 5 min of antibody circulation, excess antibody and neutrophils (those not sequestered within the pulmonary microcirculation) were flushed by cardiac perfusion with PBS during exsanguination. Finally, the lung lobes were excised and harvested, then homogenized using a lung dissociation kit (Miltenyi Biotec, USA) and a tissue homogenizer (gentleMACS^TM^ Dissociator, Miltenyi Biotec, USA). The lung homogenates were passed through 0.7 μm cell strainers, then stained in FACS buffer with PE anti-LyB.2 (to identify all neutrophils), and BV421 anti-CD45.2 (to identify donor neutrophils).

### Flow cytometry

All flow cytometry antibodies were procured from BioLegend unless otherwise stated. The antibodies used were BV421^TM^ anti-Ly6G (Clone 1A8), PE anti-Ly6G (Clone 1A8), APC-Fire^TM^ 750 anti-Ly6G (Clone 1A8), APC anti-CD63 (Clone NV-G2), PE anti-CD35 (Clone 7E9), APC-Cy7 anti-CD11b (Clone M1/70), BV421^TM^ anti-CD45.2, PE anti-LyB.2 (Clone REA115 | 7/4) (Miltenyi Biotec Inc.), Alexa Fluor^TM^ 647 anti-mouse MPO (Clone 8F4/MPO) and Alexa Fluor^TM^ 647 Mouse IgG1 κ Isotype Control (Clone MOPC-21) (BD Pharmingen^TM^). To perform flow cytometry on blood cells, RBC lysis of the blood samples was performed using RBC Lysis Buffer (BioLegend), followed by washing with FACS buffer (1%FBS/PBS), and staining with antibodies. Samples were analyzed using MACSQuant 10 or 16 (Miltenyi Biotec Inc.).

### Intracellular killing assay

*S. aureus* strain USA300-sGFP was a kind gift from Alexander Horswill (University of Colorado). *S. aureus* was grown in tryptic soy broth for 18 h at 37°C with 250 RPM shaking. Overnight cultures were subcultured to obtain bacteria in the logarithmic growth phase. Once OD600 reached 0.8, *S. aureus* was pelleted and washed twice with PBS. Then *S. aureus* was stained with ATTO647N ester (Sigma-Aldrich) at a final concentration of 0.5 μg/mL in the dark for 30 min. After 30 min, surplus dye in the solution was quenched for 3 min using assay buffer (Hank’s balanced salt solution/10% fetal bovine serum/20 mM HEPES) while also allowing for bacterial opsonization, followed by washing once with the assay buffer. Finally, the suspension was brought to OD600 of 0.5 using assay buffer and then added to 1 million bone marrow cells per sample at MOI 10 followed by incubation at 37°C. After 15 min of bacterial uptake by the neutrophils, the suspensions were washed with 200 ng/mL lysostaphin followed by one wash with assay buffer. The pellets were then resuspended in 100 μL assay buffer each. For the intracellular *S. aureus* killing assay, this was considered time zero. 20 μL from each sample was collected in FACS tubes on ice while the remaining samples were incubated for a period of 90 min at 37°C. After 90 min, samples were put on ice to stop the reaction. The samples were washed twice with ice-cold FACS buffer (1%FBS in PBS), stained with PE anti-Ly6G, BV421 anti-CD63 antibodies, and analyzed on a MACSQuant 10 or 16 flow cytometer. Data were analyzed using FlowJo software.

For intracellular staining of MPO in neutrophils infected with *S. aureus*, cells were treated with Nexinhib20 or vehicle and surface-stained with anti-Ly6G antibody simultaneously for 1 hour at room temperature, followed by washing with FACS buffer. The cells were then exposed to GFP-expressing *S. aureus* for 90 min, washed once with 200 ng/mL lysostaphin, and then washed with FACS buffer. Next, the cells were fixed and permeabilized using BD Cytofix/Cytoperm™ Fixation/Permeabilization Kit (Waters^TM^ Biosciences, Cat. No. 554714) as per the manufacturer’s protocol. Finally, the samples were intracellularly stained using anti-mouse MPO antibody or its isotype control, washed, and re-suspended in FACS buffer for flow cytometric analysis.

### ELISA

Plasma MPO was analyzed using the Mouse Myeloperoxidase ELISA kit from Abcam (Cat. 275109) as per the manufacturer’s protocol.

### Western blot

Following NP differentiation, the cells were lysed using RIPA buffer (Thermo Fisher Scientific) containing Halt^TM^ protease and phosphatase inhibitor cocktail (Thermo Fisher Scientific). Harvested proteins were resolved by SDS-PAGE and then transferred to PVDF membranes. The membranes were incubated overnight with anti-Rab27a (Cell Signaling Technology, USA) primary antibody at 4°C. After extensive washing, the blots were then incubated with HRP-conjugated secondary antibody for 1 h at room temperature, followed again by extensive washing. Finally, the blots were developed using ECL substrate (Thermo Fisher Scientific) and imaged using Chemidoc gel documentation system (BioRad).

### Statistical analysis

All statistical analyses were performed using the GraphPad Prism 10 software. Data are represented as mean±standard error of the mean (SEM). Comparisons between two groups were performed using paired or unpaired two-tailed Student’s t-tests. For comparisons among multiple groups, one-way or two-way analysis of variance (ANOVA) was performed with Tukey’s or Dunnett’s multiple comparison test, as appropriate. Data normality was assessed using the Shapiro-Wilk normality test. When the assumptions of normality were not met, the nonparametric Kruskal–Wallis test followed by Dunn’s multiple-comparisons test was used for one-way analyses. A *p* value < 0.05 was considered statistically significant.

## Acknowledgement

We are thankful to those who provided resources used in these studies, including Patrice Dubreuil (SCF-producing CHO cells), Paul Ekert (inducible HoxB8 plasmid), and Alexander Horswill (strains of USA300). Schematic diagrams were created using BioRender.com.

## Author contributions

A.B.: Designing research studies, conducting experiments, analyzing data, and writing the manuscript.

A.U., S.P., L.I.L., M.A.M., L.S.R.: assisting with experiments

C.T.L.: Supervising the project, designing research studies, analyzing data, editing the manuscript.

## Supplementary figures

**Fig S1:**
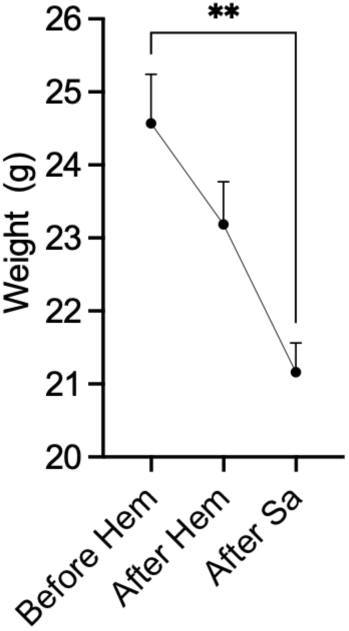
Supplementary data related to Figure 1. Weight of mice before being subjected to hemorrhage, after 24 h of hemorrhage, and after 24 h of *S. aureus* respiratory infection of those hemorrhaged mice. Data are represented as mean±SEM. Ordinary one-way ANOVA was performed with Tukey’s multiple comparison test; each group n=5 mice, **p<0.01

**Fig. S2:**
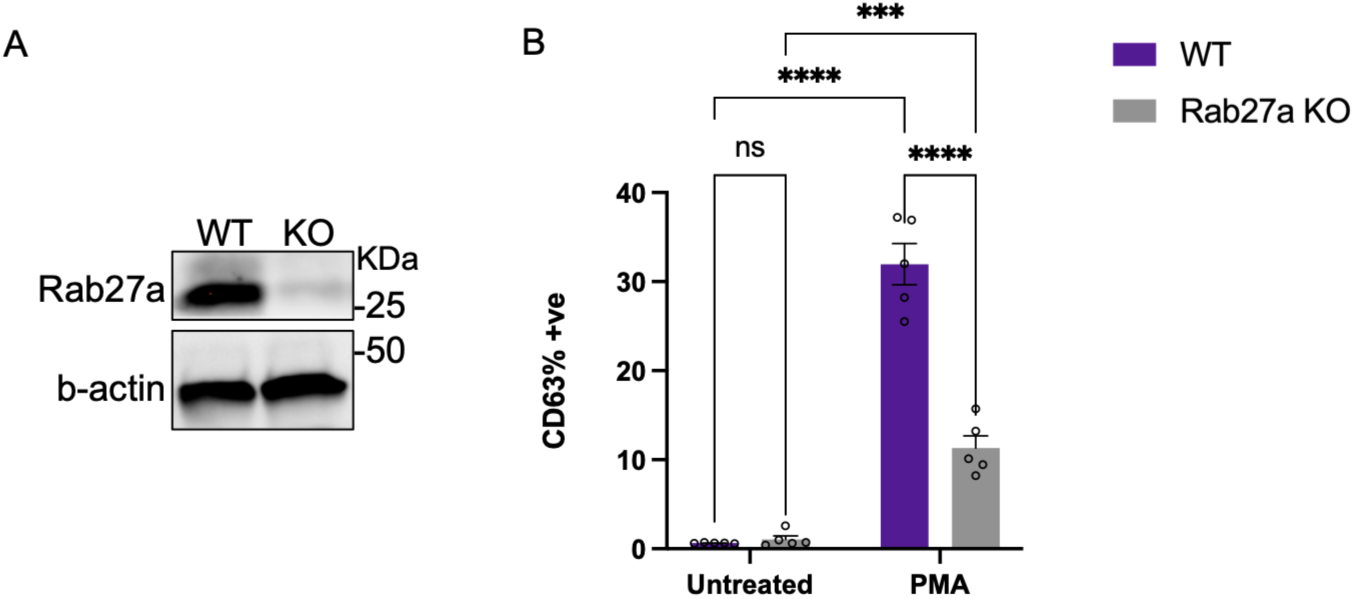
Supplemental data related to Figure 2. Rab27a KO and functional validation. **(A)** Proteins were isolated from differentiated neutrophils and subjected to Western blotting to probe for Rab27a KO using specific antibody. **(B)** Functional validation of knock-out was performed by stimulating the differentiated neutrophils with phorbol 12-myristate 13-acetate (PMA) at 100 ng/mL for 10min followed by flow cytometry analysis of surface CD63 expression as a read-out of granule exocytosis. Data are represented as mean±SEM. Two-way analysis of variance (ANOVA) was performed with Tukey’s multiple comparison test; ***p<0.001, ****p<0.0001

**Fig. S3:**
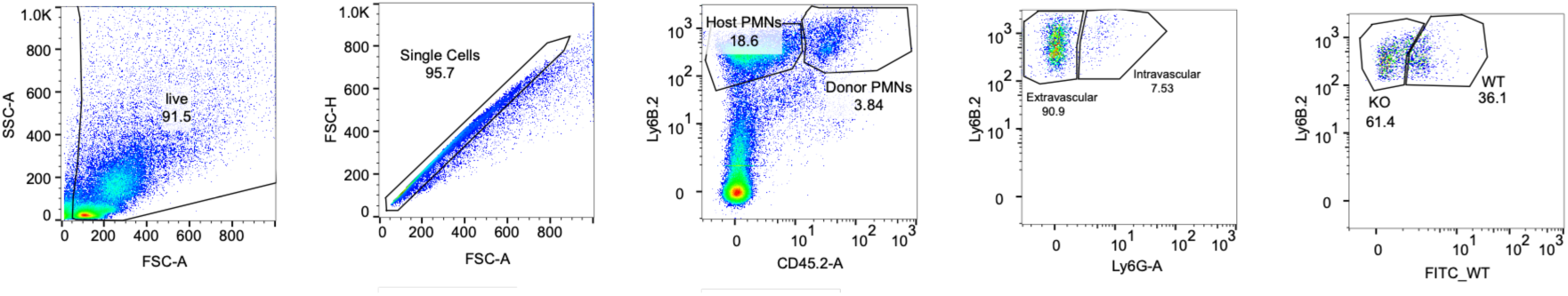
Supplemental data related to Figure 2. Gating strategy to differentiate CD45.2+ donor WT and Rab27a KO intravascular neutrophils from the extravascular neutrophils.

**Fig. S4:**
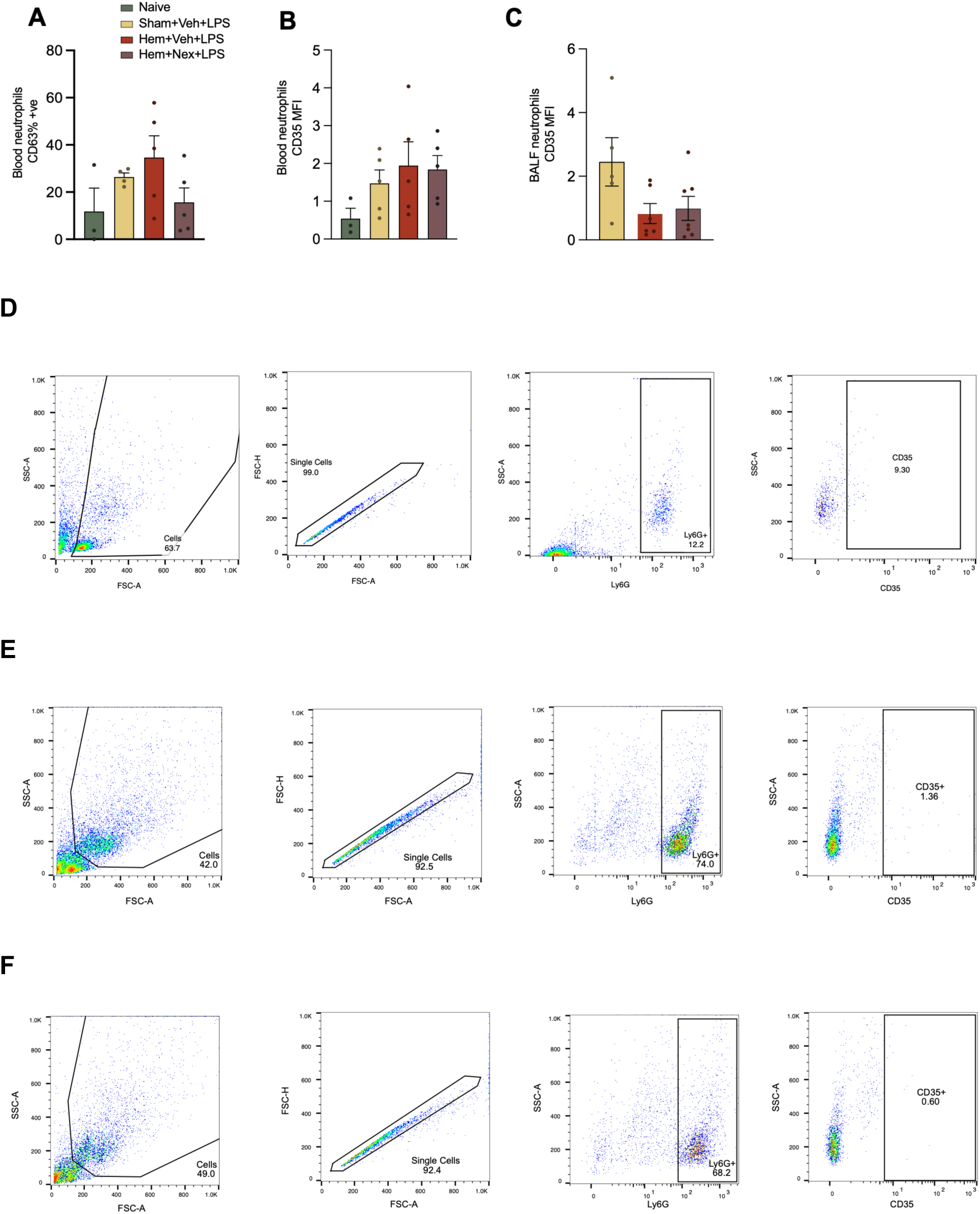
Supplemental data related to Figure 3. Analysis of surface expression of CD63 in blood neutrophils **(A)** and CD35 in blood and BALF neutrophils **(B, C)** by flow cytometry. Data normality was assessed using Shapiro-Wilk normality test. One-way ANOVA was performed with Tukey’s multiple comparison test. **(D-F)** Gating strategy for flow cytometric analysis of CD35 in blood **(D)**, BALF **(E)**, and fluorescence minus one (FMO) for CD35 **(F)**.

**Fig. S5:**
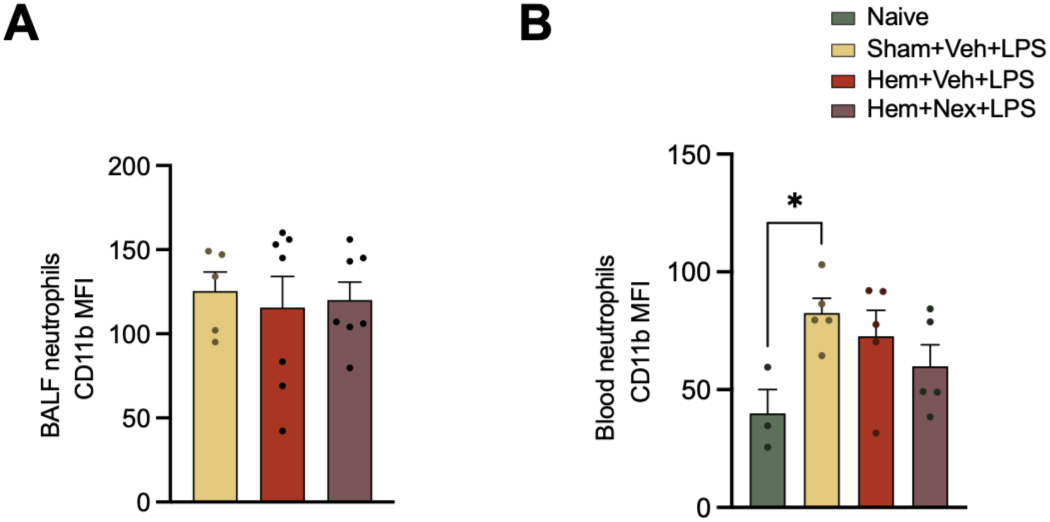
Supplemental data related to Figure 3. Analysis of surface expression of CD11b in BALF neutrophils **(A)** and blood neutrophils **(B)** by flow cytometry. Data normality was assessed using Shapiro-Wilk normality test. One-way ANOVA was performed with Tukey’s multiple comparison test; *p<0.05

